# Covalent Surface Modification Effects on Single-Walled Carbon Nanotubes for Multimodal Optical Applications

**DOI:** 10.1101/837278

**Authors:** Linda Chio, Rebecca L. Pinals, Aishwarya Murali, Natalie S. Goh, Markita P. Landry

## Abstract

Optical nanoscale technologies often implement covalent or noncovalent strategies for the modification of nanoparticles, whereby both functionalizations are leveraged for multimodal applications but can affect the intrinsic fluorescence of nanoparticles. Specifically, single-walled carbon nanotubes (SWCNTs) can enable real-time imaging and cellular delivery; however, the introduction of covalent SWCNT sidewall functionalizations often attenuates SWCNT fluorescence. Herein, we leverage recent advances in SWCNT covalent functionalization chemistries that preserve the SWCNT’s pristine graphitic lattice and intrinsic fluorescence and demonstrate that such covalently functionalized SWCNTs maintain fluorescence-based molecular recognition of neurotransmitter and protein analytes. We show that the covalently modified SWCNT nanosensor fluorescence response towards its analyte is preserved for certain nanosensors, presumably dependent on the steric hindrance introduced by the covalent functionalization that hinders noncovalent interactions with the SWCNT surface. We further demonstrate that these SWCNT nanosensors can be functionalized via their covalent handles to self-assemble on passivated microscopy slides, and discuss future use of these dual-functionalized SWCNT materials for multiplexed applications.

---

Nanomaterials are nanoscale particles that have been leveraged for biological applications such as imaging, gene or drug delivery, and therapeutics.^1,2^ Nanomaterials can offer advantages over biological materials for said applications due to their tunable physicochemical properties that enable the manipulation of nanoparticles with different chemistries to create multiple modalities on a single particle. In particular, single-walled carbon nanotubes (SWCNTs) have been used as cellular delivery vehicles, fluorescent nanosensors, and implantable diagnostics.^3–6^Covalent and noncovalent SWCNT surface modifications enable the dispersion of hydrophobic SWCNTs in aqueous solution for their use as delivery agents and as nanosensors: covalent chemical functionalization has been used to add synthetic handles to attach cargo useful for molecular recognition and targeted delivery,^7–9^ whereas noncovalent chemical functionalization is a preferred approach for sensing applications to preserve the SWCNT’s intrinsic fluorescent properties. As with many other classes of nanoparticles, chemical functionalization of the SWCNT surface will often affect the nanoparticle’s various optical, physical, and material properties.^10^

SWCNTs are an attractive class of nanomaterial for sensing applications, owing to the unique near-infrared (NIR) fluorescence of SWCNTs in the short wavelength infrared range of ∼1000-1300 nm. This region of NIR fluorescence is optimal for biological imaging, as the long wavelength light is minimally attenuated in biological tissues via reduced scattering and absorption of photons.^11,12^ For SWCNT use as molecular nanosensors, it is necessary to preserve the intrinsic NIR fluorescence that arises from Van Hove transitions resulting from the density of states of the SWCNT lattice.^13^ This fluorescence has been employed to develop a class of nanosensors that undergo fluorescence modulation upon selective binding of bioanalytes such as neurotransmitters, reactive nitrogen species, metabolites, peptides, and proteins.^14–19^ The selectivity of these varied SWCNT nanosensors is created through a phenomenon termed corona phase molecular recognition (CoPhMoRe). CoPhMoRe nanosensors are capable of binding specific analytes through a constrained surface-adsorbed state formed by the noncovalent association of the SWCNT surface and a coating, such as polymers or phospholipid, not necessarily known to bind the analyte of interest. While SWCNT-based nanosensors generated with CoPhMoRe have shown recent success for imaging analytes *in vivo*^6,20,21^ and for *ex vivo* imaging neuromodulation in acute brain slices,^22^ their use has involved undirected biodistribution of SWCNTs in the tissue under investigation. For the purposes of tissue-specific or targeted sensing, inclusion of targeting moieties such as aptamers or proteins can be achieved via direct covalent attachment to SWCNT surfaces. Covalent chemistries offers several advantages over noncovalent chemistries for this attachment such as the formation of a strong covalent bond between the SWCNT surface and the targeting moiety and the preservation of the targeting moiety’s structure through the availability of biocompatible bioconjugation attachment chemistries.^23^ However, given that covalent modifications to SWCNTs often compromise intrinsic fluorescence, to date, simultaneous covalent and noncovalent functionalization of the SWCNT surface for sensing applications remains an outstanding challenge.

Obstacles for simultaneous covalent and noncovalent functionalization of fluorescent SWCNTs arise from surface defects created by covalent reactions. When the sp^2^-hybridized carbon lattice of the SWCNT surface is disrupted via covalent modification, often non-radiative exciton recombination predominates, attenuating or obliterating SWCNT fluorescence which compromises the fluorescent readout of the SWCNT-based optical nanosensor.^24^ Mild covalent SWCNT modifications via end-cap or defect engineering enable the creation of controlled sp^3^ defects that modulate the SWCNT bandgap, resulting in a defect-based fluorescence red shift in the SWCNT fluorescence.^25–28^ Additionally, recent developments in SWCNT chemistry have established a covalent functionalization reaction that re-aromatizes defect sites to re-form the original, pristine SWCNT lattice and restore intrinsic fluorescence.^29^ This development could enable synergistic combination of covalent and noncovalent functionalization strategies to confer multiple functionalities to SWCNT-based technologies, such as theranostics, targeted fluorescence imaging, and towards understanding the fate of functionalized SWCNTs upon cellular delivery.

Herein, we combine covalent and noncovalent SWCNT functionalizations to study the effects of covalent SWCNT modification on CoPhMoRe-based SWCNT nanosensors that rely on noncovalent functionalization for sensing. We synthesized and characterized several SWCNTs with different covalent surface modifications that are subsequently functionalized with noncovalent coatings for downstream use as fluorescent nanosensors. We show how the addition of surface groups impacts both the fluorescence and analyte responsivity of certain CoPhMoRe-based SWCNT nanosensors. We further demonstrate how the addition of charged chemical functionalizations affects the corona formation and overall SWCNT assembly stability. Finally, we combine these findings to create dual-functional SWCNTs with targeted recognition and fluorescence sensing capabilities.

## Results and Discussion

We generated defect-free covalently functionalized SWCNTs as previously reported^29,30^ by performing a chemical re-aromatization reaction using unfunctionalized SWCNTs (pristine-SWCNTs) and cyanuric chloride to produce triazine-functionalized SWCNTs at low and high-labeling densities, denoted Trz-L-SWCNTs and Trz-H-SWCNTs, respectively. Trz-H-SWCNTs were further functionalized through nucleophilic substitution of the chlorine on the triazine with a primary amine to create a library of surface-functionalized SWCNTs at positions denoted by variable R groups (Fig. 1a and 1b). In this manner, we reacted Trz-H-SWCNTs with cysteine to form thiol-functionalized SWCNTs (SH-SWCNTs).

**Figure 1.**
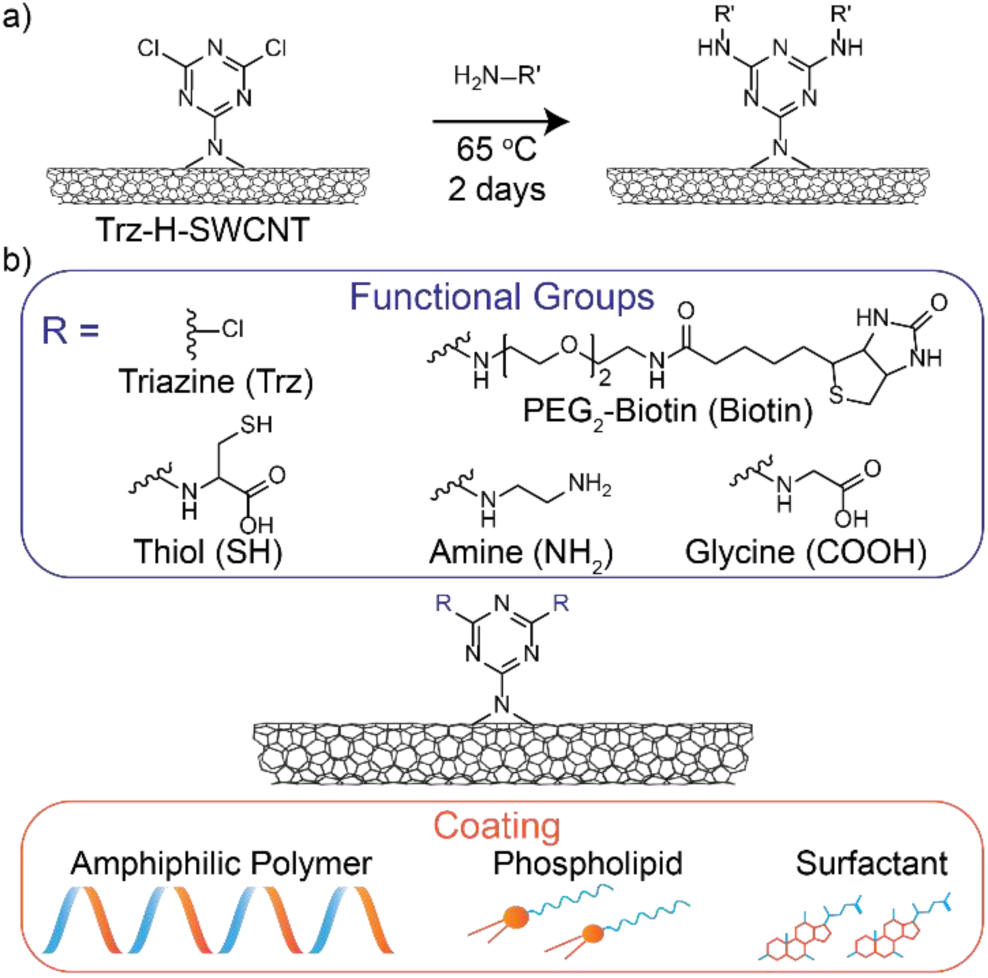
Functionalization of SWCNTs for nanosensor generation. a) Functional groups are added to Trz-H-SWCNT via a nucleophilic substitution reaction whereby primary amines replace the chlorines of the triazine group. b) Overview of the different functional groups and SWCNT coatings tested to create multifunctional fluorescent nanosensors.

We next assessed the impact of the triazine and thiol covalent SWCNT surface functional groups on the performance of CoPhMoRe nanosensors generated from Trz-L-SWCNTs, Trz-H-SWCNTs and SH-SWCNTs (Fig. 2a). We tested several previously reported SWCNT-based nanosensors to image dopamine, fibrinogen, and insulin analytes. Specifically, when noncovalently adsorbed to the SWCNT surface, single-stranded DNA (ssDNA) oligomers (GT) _15_ and (GT)_6_ form known nanosensors for dopamine,^14,31^ DPPE-PEG5K phospholipid for fibrinogen,^17^ and C_16_-PEG2k-ceramide phospholipid for insulin.^18^ To generate each nanosensor, we induced noncovalent association of the SWCNTs with each coating through π-π aromatic stabilization and hydrophobic packing via probe-tip sonication or dialysis as previously established (see methods for details). We prepared dopamine, fibrinogen, and insulin nanosensors with both pristine and functionalized SWCNTs. We expand upon the findings of previous literature that covalently-functionalized Trz-L-SWCNTs, Trz-H-SWCNTs, and SH-SWCNTs maintain their intrinsic optical properties and show that this remains the case when dispersed with their respective surfactant, phospholipid, or polymer coatings (Fig. S1, Fig. S2), prior to assessing their use as fluorescent optical nanosensors.

**Figure 2:**
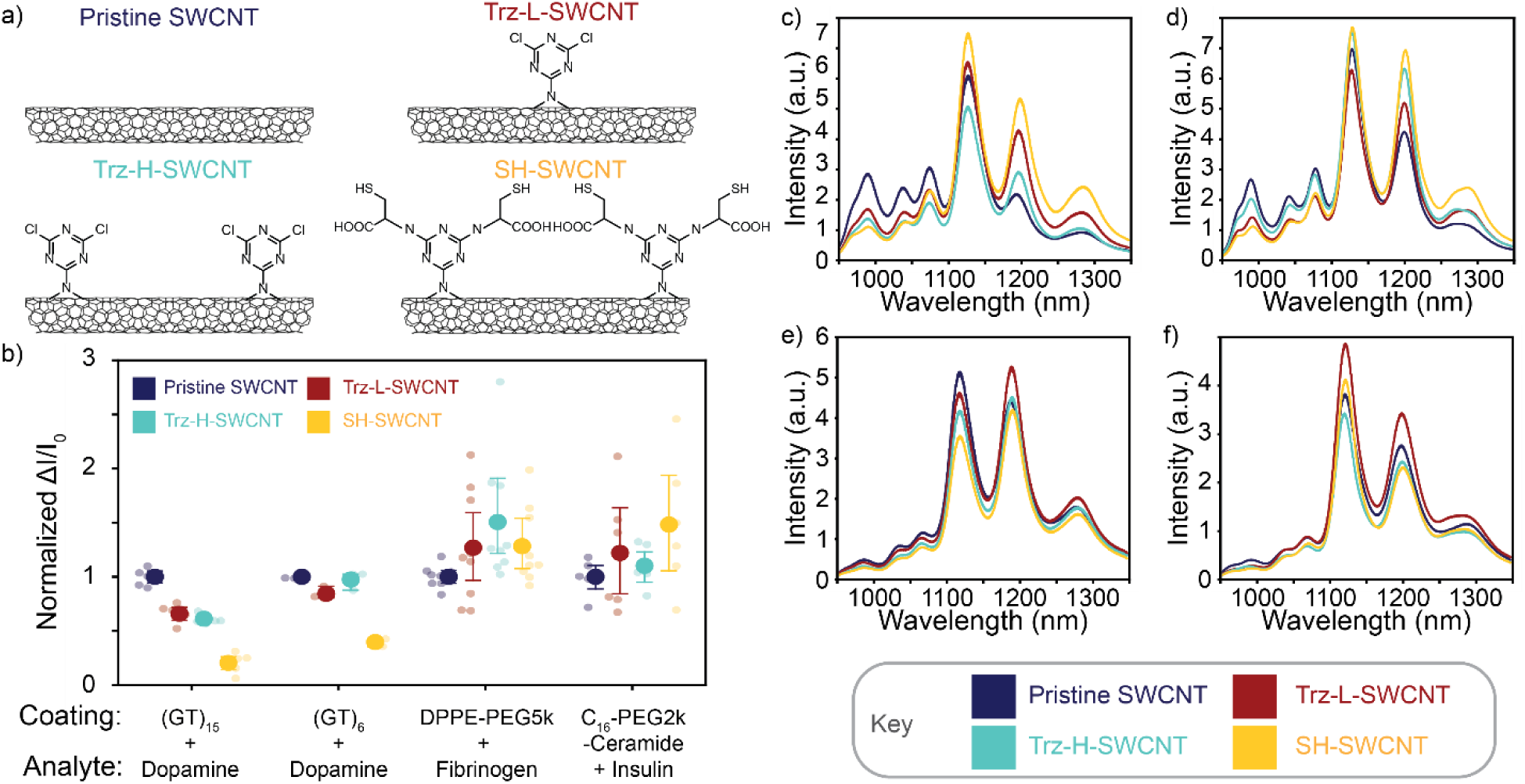
Effect of covalent and noncovalent functionalization on SWCNT nanosensor fluorescence response. a) Structures of covalently functionalized SWCNTs investigated in this study. b) Normalized fluorescence change ΔI/I_0_ of 5 mg/L covalently modified SWCNT nanosensors upon addition of their respective dopamine (100 μM), fibrinogen (1 mg/mL), and insulin (20 μg/mL) analytes, normalized to the response of nanosensors generated from pristine-SWCNT (large dots denote mean ΔI/I_0_, each small faded dot denotes an experimental replicate, and error bars denote standard deviation for n = 3 to 9 trials). Covalently functionalized ssDNA-SWCNT nanosensors sensitive to the coating’s structural conformation show an attenuated response to analyte, as compared to structure-independent phospholipid coatings. Fluorescence spectra of concentration-normalized samples of 5 mg/L c) (GT)_15_-SWCNT, d) (GT)_6_-SWCNT, e) DPPE-PEG5k-SWCNT, or f) C_16_-PEG2k-Ceramide-SWCNT.

By comparing the performance of nanosensors generated from pristine versus functionalized SWCNTs, we quantified functionalization-dependent fluorescence performance of nanosensors upon exposure to their respective analytes of dopamine, fibrinogen, and insulin (Fig. 2b). We measured the fluorescence change of 5 mg/L phospholipid or polymer suspended pristine-SWCNT, Trz-L-SWCNT, Trz-H-SWCNT, and SH-SWCNT nanosensors upon exposure to their respective analytes. Responses were measured before and 30 minutes after the addition of 100 μM dopamine to (GT)_15_- and (GT)_6_-SWCNT nanosensors, 1 mg/mL of fibrinogen to DPPE-PEG5k-SWCNT nanosensors, and 20 μg/mL insulin to C_16_-PEG2k-ceramide-SWCNT nanosensors. Changes in fluorescence were calculated and normalized to the changes measured in pristine-SWCNT nanosensors to determine how surface functionalization impacts CoPhMoRe sensing.

The performance of (GT)_15_ dopamine nanosensors decreased when covalently functionalized SWCNTs were used as compared to pristine-SWCNTs. We calculated the normalized nanosensor fluorescence response, ΔI/I_0_, for (GT)_15_-Trz-L-SWCNTs (0.661 ± 0.118, mean ± SD), (GT)_15_-Trz-H-SWCNTs (0.613 ± 0.083, mean ± SD), and (GT)_15_-SH-SWCNTs (0.207 ± 0.095, mean ± SD), compared to a nanosensor fluorescence response for the original dopamine nanosensor constructed from (GT)_15_-pristine-SWCNTs (1.0 ± 0.121, mean ± SD). Previous molecular simulations postulate that the longer polymer (GT)_15_ forms a helical conformation on the surface of the nanotube, while the shorter polymer (GT)_6_ forms rings on the surface.^32^ To assess the impact of ssDNA polymer length on adsorption to a covalently functionalized SWCNT surface, we tested the dopamine responsivity of of Trz-L-SWCNTs, Trz-H-SWCNTs, and SH-SWCNTs noncovalently functionalized with (GT)_6_ versus (GT)_15_ ssDNA. We hypothesize that the corona adopted by the ring-forming (GT)_6_ is less sterically perturbed by the addition of chemical functional groups on the SWCNT surface. Unlike dopamine nanosensors generated from (GT) _15_, we found that the normalized nanosensor fluorescence responses for (GT)_6_ suspended Trz-L-SWCNTs, Trz-H-SWCNTs, and SH-SWCNTs, exhibited a recovery of fluorescence response: ΔI/I_0_ for (GT)_6_-Trz-L-SWCNTs (0.844 ± 0.051, mean ± SD), (GT)_6_-Trz-H-SWCNTs (0.974 ± 0.073, mean ± SD), and (GT)_6_-SH-SWCNTs (0.396 ± 0.031, mean ± SD), are all closer to the fluorescence response for the dopamine nanosensor constructed from (GT)_6_-pristine-SWCNTs (1.0 ± 0.026, mean ± SD) compared to functionalized SWCNT dopamine sensors generated with (GT) _15_. The normalized ΔI/I_0_ performance of (GT)_6_-based dopamine nanosensors represent a 1.27, 1.59, and 1.91-fold increase in performance over (GT)_15_-based dopamine nanosensors for nanosensors generated from Trz-L-SWCNTs, Trz-H-SWCNTs, and SH-SWCNTs, respectively, suggesting that shorter ring-forming ssDNA oligomers are less sterically hindered by covalent SWCNT surface modifications than longer helix-forming ssDNA oligomers. (GT)_6_-SH-SWCNT nanosensors still exhibit a greatly attenuated fluorescence response to dopamine, which could be due to the bulkier thiol functional groups as compared to the chlorine of the triazine SWCNTs. We further confirmed that differences in nanosensor performance are not due to intrinsic differences in baseline fluorescence of the concentration-normalized SWCNT samples and can therefore be attributed to surface steric hinderance or the intrinsic chemical properties of the functional groups (Fig. 2 c-f).

To further probe the effect of steric contributions to noncovalent CoPhMoRe corona formation on the SWCNT surface, we tested the fluorescence response of DPPE-PEG5k-SWCNT fibrinogen nanosensors. We expect that the adsorption of DPPE-PEG5k phospholipids, which adopt a self-assembled membrane-like structure on SWCNT surfaces, will be relatively sterically unhindered when binding to covalently-functionalized SWCNTs.^33,34^ Upon addition of 1 mg/mL fibrinogen, we observed equivalent fluorescence responses of fibrinogen based on covalently-functionalized SWCNT nanosensors relative to those made from pristine-SWCNTs, corroborating that steric effects contribute to nanosensor attenuation when SWCNTs are suspended with conformationally-dependent amphiphilic polymers such as ssDNA. DPPE-PEG5k coated Trz-L-SWCNTs (1.268 ± 0.514, mean ± SD), Trz-H-SWCNTs (1.506 ± 0.575, mean ± SD), and SH-SWCNTs (1.282 ± 0.358, mean ± SD) all maintained their response to fibrinogen, compared to the original fibrinogen nanosensor constructed from DPPE-PEG5k coated pristine-SWCNTs (1.0 ± 0.097, mean ± SD). We also tested another phospholipid-based CoPhMoRe nanosensor, C_16_-PEG2k-ceramide-SWCNT, which responds to insulin. Similarly, C_16_-PEG2k-ceramide coated Trz-L-SWCNTs (1.220 ± 0.561, mean ± SD), Trz-H-SWCNTs (1.101 ± 0.186, mean ± SD), and SH-SWCNTs (1.482 ± 0.617, mean ± SD) all maintained their response to insulin, compared to a nanosensor fluorescence response for the native insulin nanosensor constructed from pristine-SWCNTs (1.0 ± 0.157, mean ± SD). We hypothesize that phospholipid-based nanosensor responses are maintained with covalently functionalized SWCNTs because phospholipids are smaller molecules than amphiphilic polymers and may pack more densely on the surface of the SWCNT, maintaining a corona similar to that formed on pristine-SWCNTs.

We next investigated how the intrinsic properties of the covalent functionalization, such as charge, affect the formation of noncovalent coatings on the SWCNT surface. We generated SWCNTs with a positive charge through covalent addition of ethylenediamine (NH_2_-SWCNTs) and, separately, SWCNTs with a negative charge through covalent addition of glycine (COOH-SWCNTs) to the triazine handles of Trz-H-SWCNTs (Fig. 3a). We confirmed the generation of these charged SWCNT constructs through X-ray photoelectron spectroscopy and Fourier-transform infrared spectroscopy (Fig. S3 and S4). We sought to assess whether the (GT)_15_-– SWCNT dopamine nanosensor yields, as a proxy for coating stability, would be affected by the surface charges using a methanol-driven coating exchange method.^35,36^ We show that the yield of (GT)_15_-coated SWCNTs is highest for the positively charged NH_2_-SWCNTs (Fig. 3b), as expected given the negative charge of (GT)15 (Fig. S5). The positive charge of the amine group on the surface of NH_2_-SWCNTs presumably favors the association with negatively charged (GT)_15_, and displays a 22.9% nanosensor yield that is significantly higher than those of pristine-SWCNTs and COOH-SWCNTs (p < 0.05, uncorrelated independent student T-test). The yields for (GT)_15_ coated pristine-SWCNTs and COOH-SWCNTs were less than 10% and not significantly different (p-value = 0.47 > 0.05, uncorrelated independent student T-test), suggesting (GT)_15_ is more stably adsorbed to NH_2_-SWCNTs than to COOH- or pristine-SWCNTs. We attribute the lower ssDNA-COOH-SWCNT yield to the negatively charged COOH-SWCNT, which electrostatically repels the negative (GT)_15_-polymer. These results indicate that coating formation could be driven or hindered by the intrinsic charge properties of the covalent functional group.

**Figure 3:**
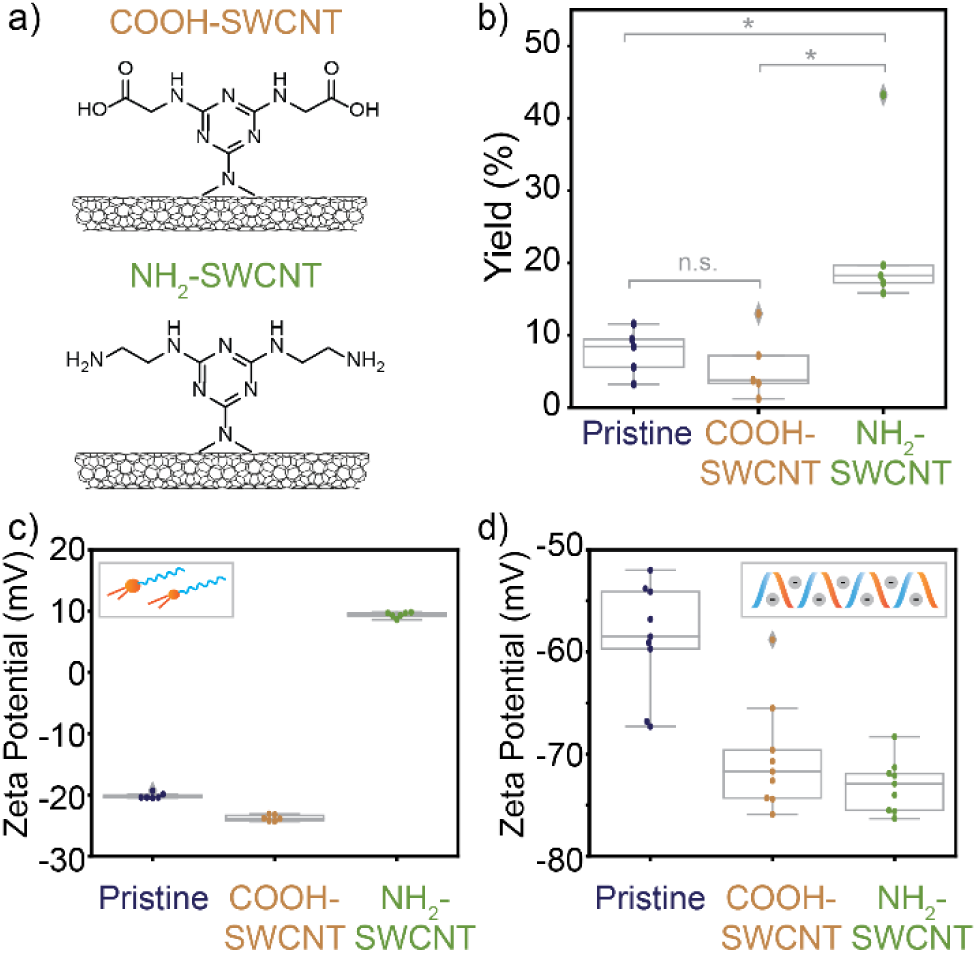
SWCNT surface charge imparted by covalent functionalization impacts subsequent noncovalent functionalization. a) Structures of negatively-charged COOH-SWCNT and positively-charged NH_2_-SWCNT. b) Box-and-whisker plot showing percent yields of negatively-charged (GT)_15_-coated SWCNTs upon methanol exchange. (n = 5 trials and * denotes p < 0.05 (uncorrelated independent student T-test)). Box-whisker-plots of zeta potential measurements (n = 6) of SWCNTs with c) a neutral C_16_-PEG2k-ceramide phospholipid coating or d) negatively-charged (GT)_15_ ssDNA polymer coating.

To demonstrate that the charge properties of NH_2_ and COOH covalently functionalized SWCNTs can be maintained, we suspended COOH-SWCNTs, NH_2_-SWCNTs, and pristine-SWCNTs with a neutral (uncharged) phospholipid, C_16_-PEG2k-ceramide. The zeta potentials, which characterize the electrokinetic potential at the slipping plane and provide a proxy for particle surface charge, were measured for these three SWCNT constructs with C_16_-PEG2k-ceramide coatings. C_16_-PEG2k-ceramide coated COOH-SWCNTs (−23.8 mV ± 0.501, mean ± SD) displayed the most negative zeta potential and C_16_-PEG2k-ceramide coated NH_2_-SWCNTs (9.38 mV ± 0.446, mean ± SD) displayed the most positive zeta potential, as compared to C_16_-PEG2k-ceramide coated pristine-SWCNTs (−20.1 mV ± 0.468, mean ± SD) (Fig. 3c, Fig. S6a). We repeated this experiment with a negatively charged (GT)_15_ polymer coating instead, and measured the zeta potentials for pristine-SWCNTs (−55.0 mV ± 3.69, mean ± SD), COOH-SWCNTs (−70.39 mV ± 5.33, mean ± SD), and NH_2_-SWCNTs (−73.1 mV ± 2.54, mean ± SD). As expected, all complexes showed a more negative surface charge than the complexes formed with the C_16_-PEG2k-ceramide coating (Fig. 3d, Fig. S6b). Tuning of intrinsic properties of SWCNT surfaces in this manner could aid SWCNT design for cellular delivery applications, where charge is a driving factor for nanoparticle internalization and subsequent cytotoxicity.^37^

Finally, we tested the use of covalent modifications to provide additional function to optical SWCNT nanosensors. Covalent attachment offers a method for the addition of molecular recognition elements such as antibodies and nanobodies, targeting modalities, and therapeutics.^6,38^ We explored this dual functionality of SWCNTs through the attachment of biotin as an affinity pair with avidin protein as a model system (Fig. 4a). Biotin is known to form a high affinity noncovalent association with the avidin protein and its analogues such as neutravidin and streptavidin.^39^ We generated biotin-SWCNTs by covalently attaching amine-PEG2k-biotin to SWCNTs through the triazine handles of Trz-H-SWCNTs (Fig. 1a). Biotin-SWCNTs were then coated with (GT)_15_ ssDNA polymers to test the use of multifunctional SWCNTs.

**Figure 4.**
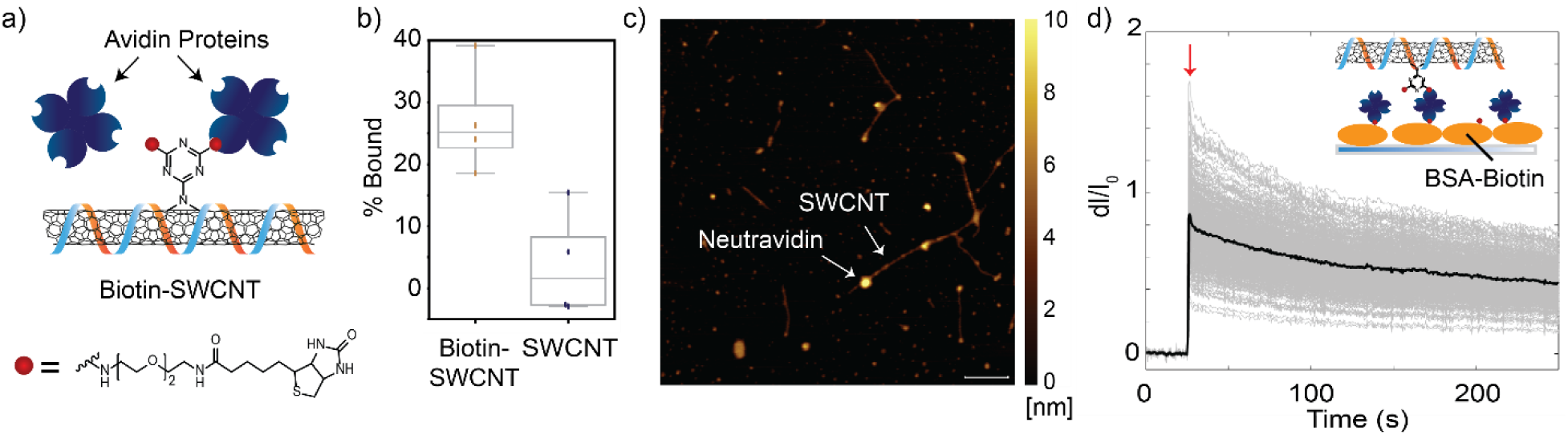
Covalent modification of SWCNTs with amine-PEG2-biotin adds functional handles for avidin protein attachment. a) (GT)_15_-biotin-SWCNTs bind tetrameric avidin proteins such as neutravidin and streptavidin. b) A mass balance shows a higher percentage of bound (GT)_15_-biotin-SWCNTs on streptavidin beads compared to (GT)_15_-pristine-SWCNTs alone. c) AFM images show neutravidin protein bound to (GT)_15_-biotin-SWCNTs. d) (GT)_15_-biotin-SWCNTs immobilized on a neutravidin-coated microscopy surface (inset) and imaged with a near-infrared epifluorescence microscope with 721 nm excitation shows fluorescence response following exposure to 25 μM dopamine (addition denoted by the red arrow). The black line denotes the average fluorescence trace with individual microscopic regions of interest denoted by the gray lines

We performed an affinity assay with streptavidin-coated magnetic beads to verify the attachment of biotin to the SWCNT surface. A significantly higher amount of (GT)_15_-biotin-SWCNTs remained bound to magnetic beads (23.0% ± 4.0, mean ± SD) than pristine-SWCNTs (0.1 % ± 5.0, mean ± SD) (Fig. 4b). The attachment of biotin to SWCNTs was further validated through atomic force microscopy (AFM). AFM was used to visualize the presence of neutravidin protein along the length of individual (GT)_15_-biotin-SWCNTs (Fig. 4c). SWCNT heights were measured as 0.5 nm to 1 nm and the neutravidin protein heights were measured as ∼4 – 6 nm similar to previous reports.^19,40^ We found that 58.9% of (GT)_15_-biotin-SWCNTs (66 out of 112 SWCNTs total from multiple fields of view) as compared to 38.8% (GT)_15_-pristine-SWCNTs (19 out of 49 SWCNTs total from multiple fields of view) appear to bind neutravidin protein (Fig. 4d). Lastly, we tested the ability to use (GT)_15_-biotin-SWCNTs for dopamine imaging. (GT)_15_-biotin-SWCNTs were deposited onto a microscope slide surface-functionalized with a neutravidin monolayer, then the surface-bound nanosensors were exposed to 25 μM dopamine. As shown in Figure 4d, exposure to dopamine generated an integrated fluorescence response ΔI/I_0_ = 0.8658 ± 0.1986 (mean ± SD), suggesting that SWCNTs that are dual-functionalized with both covalent handles and noncovalent polymeric ligands can be used for analyte-specific imaging applications. Taken together, these results suggest the potential of using covalent functional groups on SWCNTs for multifunctional applications.

## Conclusions

We generated and characterized the properties of optically-active, covalently functionalized SWCNTs for use as fluorescent nanosensors. Comparisons of covalently functionalized SWCNT to pristine-SWCNT showed that the introduction of chemical handles could impact a SWCNT-based nanosensor response to its analyte, depending on the structural perturbation of the noncovalent coating adsorption upon corona formation. Nanosensors generated with long amphiphilic polymers showed the greatest analyte-specific fluorescence response attenuation following covalent addition of chemical handles on the SWCNT surface, whereas nanosensors generated from phospholipid-based coronas did not exhibit analyte-specific fluorescence response attenuation. Notably, following covalent functionalization of the SWCNT, all SWCNT complexes successfully formed through the noncovalent association of amphiphilic polymers or phospholipids, and were still capable of acting as optical nanosensors. These results validate the use of covalently functionalized SWCNTs as CoPhMoRe fluorescent nanosensors with unique analyte specificity, corona stability, and spatiotemporal readout appropriate for *in vivo* and *ex vivo* imaging.

This study further suggests the promise of utilizing dual covalent and noncovalent SWCNT functionalizations to create multifunctional nanoscale tools. The surface charge of SWCNT complexes can be altered via addition of covalent handles without perturbation of the SWCNT fluorescence, enabling CoPhMoRe-based sensing together with chemical handles for charge-based functionalizations and addition of ligands based on electrostatics. Furthermore, we show successful covalent attachment of the targeting moiety biotin to (GT)_15_-SWCNT dopamine nanosensors and show the biotin group can be used for surface-immobilized dopamine imaging with near-infrared microscopy. These results demonstrate the possibility of dual covalent and noncovalent SWCNT functionalization for multiplexed purposes with future applications in targeted delivery, theranostics, targeted fluorescence imaging, and towards understanding the fate of functionalized SWCNTs upon cellular delivery.

## Methods

### Materials

All chemicals unless otherwise noted were purchased from Sigma-Aldrich. Raw high pressure carbon monoxide (HiPCO) synthesized single-walled carbon nanotubes (SWCNTs) were purchased from NanoIntegris. EZ-Link amine-PEG2-biotin and neutravidin protein was purchased from ThermoFisher Scientific. Streptavidin magnetic beads were purchased from New England BioLabs.

### Synthesis of Triazine-Functionalized SWCNTs (Trz-H-SWCNTs and Trz-L-SWCNTs)

Synthesis of Trz-SWCNTs was adapted from reference 29. In brief, SWCNTs (1 g) were dispersed in N-methyl-2-pyrrolidone (150 ml) in a round bottom flask and bath sonicated for 1 hr. The dispersion was stirred for 1 hr at 25 °C and cooled down to 0 °C. 2,4,6-1,3,5-trichlorotriazine (10 g, 54 mmol) was dissolved in N-methyl-2-pyrrolidone (50 ml) and the obtained solution was slowly added to the SWCNT dispersion at 0 °C. Sodium azide (1.76 g, 27 mmol) was added to the mixture and stirred for 2 hr at 0 °C followed by 12 hr stirring at either 70 °C or 25 °C to yield Trz-H-SWCNTs or Trz-L-SWCNTs, respectively. The product was purified by centrifugation and washed by re-dispersion in water and organic solvents (acetone, toluene, then chloroform), and lyophilized for storage and characterization.

### Synthesis of SH-SWCNTs or Charged SWCNTs (NH_2_-SWCNTs and COOH-SWCNTs)

Trz-H-SWCNTs (10 mg) were dispersed in dimethylformamide (DMF) (5 ml) and bath sonicated for 15 min at room temperature. Next, 100 mg of either cysteine, ethylenediamine, or glycine (for SH-SWCNTs, NH_2_-SWCNTs, and COOH-SWCNTs, respectively) and a 1.5 molar excess of triethylamine to chemical were added to the mixture that was stirred at 65 °C for 2 days. The product was purified by centrifugation and re-dispersion in two washes of 4 mL of DMF followed by two washes of 4 mL of water. SH-SWCNTs were dialyzed against water using a Slide-A-Lyzer G2 10 kDa molecular cutoff dialysis cassette (Thermo Scientific) with daily water changes for 1 week. The product was lyophilized for storage and characterization.

### Synthesis of Biotin-SWCNTs

Trz-H-SWCNTs (5 mg) were dispersed in DMF (2 mL) and bath sonicated for 15 min at room temperature. A solution of 25 mg EZ-Link amine-PEG2-biotin in 1 mL DMF was made up. 500 μL of the amine-PEG_2_-biotin solution and 86 μL of triethylamine were added to the Trz-SWCNT solution. The mixture was stirred at 65 °C for 2 days. The product was purified by centrifugation and re-dispersion in two washes of 2 mL of DMF followed by two washes of 2 mL of water. The product was then dialyzed against water using a Slide-A-Lyzer G2 10 kDa molecular cutoff dialysis cassette (Thermo Scientific) with daily water changes for 1 week. The product was pelleted by centrifugation and lyophilized for storage and characterization.

### Noncovalent Adsorption of Polymer and DPPE-PEG5k Coatings to SWCNT by Probe-Tip Sonication

(GT)_15_, (GT)_6_, and DPPE-PEG5k SWCNT sensors were constructed through probe-tip sonication. 1 mg of SWCNTs were added to 500 μL of 1× phosphate buffered saline (PBS, pH 7.4) and 1 mg of polymer or DPPE-PEG5k. The solution was bath sonicated for 10 minutes, and then probe-tip sonicated using a Cole Parmer ultrasonic processor and a 3 mm stepped microtip probe with pulses of 3-7 watts every 3 seconds for 15 minutes. The solution was subsequently allowed to equilibrate on the bench at room temperature for 1 hour before centrifugation at 16.1 × 10^3^ Relative Centrifugal Force (RCF) for 30 minutes to remove any unsuspended nanotube aggregates. Sensors formed a homogeneous dark-gray solution and concentration was characterized by UV-Vis-IR absorbance using a Shimadzu UV-3600 Plus. SWCNT concentration was calculated from absorbance at 632 nm using Beer-Lambert law with extinction coefficient, ε_632_ = 0.036 L mg^-1^ cm^-1^.^12^

### Noncovalent Adsorption of Sodium Cholate to SWCNT (SC-SWCNT)

3 mg of SWCNTs were added to 3 mL of 1 wt % sodium cholate (SC). The solution was bath sonicated for 10 minutes, and then probe-tip sonicated using a 500 W Cole Parmer Ultrasonic Homogenizer and a 6 mm stepped microtip probe at 10% amplitude every 2 seconds for 60 minutes. The solution was subsequently allowed to equilibrate at room temperature for 1 hour before centrifugation at 23.1 × 10^3^ Relative Centrifugal Force (RCF) for 50 minutes to remove any unsuspended nanotube aggregates. Assemblies formed a homogeneous dark-gray solution and concentration was characterized by UV-Vis-IR absorbance using a Shimadzu UV-3600 Plus. SWCNT concentration was calculated from absorbance at 632 nm using Beer-Lambert law with extinction coefficient, ε_632_ = 0.036 L mg^-1^ cm^-1^.^12^

### Noncovalent Adsorption of C16-PEG2k-Ceramide Coating to SWCNT through Dialysis

A solution of 25 mg/L SC-SWCNTs and 2 mg/mL C16-PEG2k-ceramide in 1% sodium cholate was added to a 1 kDa molecular weight cutoff dialysis cartridge (GE Healthcare). The solution was dialyzed against water with multiple water exchanges for a week, such that the C16-PEG2k-ceramide exchanged the sodium cholate coating by adsorbing onto the nanotube surface and replacing the small surfactant molecules.

### Near Infrared Spectroscopy of SWCNT Nanoensors

All SWCNT nanosensor solutions were diluted to a final SWCNT concentration of 5 mg/L in 1x PBS. Spectroscopic analysis was performed by measuring the resulting SWCNT photoluminescence with a home-built near infrared fluorescence microscope. Briefly, a Zeiss AxioVision inverted microscope was coupled to a Princeton Instruments IsoPlane 320 containing a liquid nitrogen-cooled Princeton Instruments PyLoN-IR 1D InGaAs array. SWCNT sensors were illuminated by a 500 mW, 721 nm laser. The spectra of SWCNT sensors were acquired, each in a separate well of a glass-bottom 384-well plate (Corning).

### X-ray Photoelectron Spectroscopy (XPS)

Samples were drop cast onto the surface of a clean silicon wafer. XPS spectra were collected with a PHI 5600/ESCA system equipped with a monochromatic Al Kα radiation source (hν = 1486.6 eV). High-resolution XPS spectra were deconvoluted with MultiPak software (Physical Electronics) by centering the C-C peak to 284.5 eV, constraining peak centers to ±0.1 eV the peak positions reported in previous literature^41^, constraining full width at half maxima (FWHM) ≤1.5 eV, and applying Gaussian-Lorentzian curve fits with the Shirley background.

### Fourier Transform Infrared Spectroscopy

Infrared spectra were measured on a Perkin-Elmer Spectrum 100 Optica FTIR spectrometer equipped with an attenuated total reflectance (ATR) accessory. 1 mg of purified and vacuum-dried SWCNT samples was deposited to the surface of the ATR and scanned.

### Methanol Exchange of Phospholipid to DNA Coating

Methanol-driven exchange of phospholipid to DNA coating was performed to be able to quantitively measure starting and final quantities of SWCNTs by absorbance spectroscopy. The following protocol has been adapted from previous studies.^35,36^ A 150 μL solution of 100 mg/L of C16-PEG2k-Ceramide-coated SWCNTs in 1× PBS (15 ng of SWCNTs) was mixed with a solution of 1 mg of DNA in 100 μL 1× PBS. Methanol was added in increments of 90 μL followed by 5 minutes of bath sonication. 7 total additions of methanol were added to the solution for a total of 630 μL of methanol. 300 μL of isopropanol was added immediately followed by brief centrifugation (17000 RCF for ∼1 s) to immediately cause the precipitation of DNA-wrapped SWCNTs in a soft pellet. The supernatant was placed in another microcentrifuge tube and centrifuged at 17000 RCF for 15 minutes to precipitate remaining DNA. The DNA pellet was resuspended in 400 μL of PBS and subsequently used to resuspend the DNA-SWCNT pellet. The solution was bath sonicated for 10 minutes, and then probe-tip sonicated using a Cole Parmer ultrasonic processor and a 3 mm stepped microtip probe with pulses of 3-7 watts every 3 seconds for 15 minutes. The solution was subsequently allowed to equilibrate at the bench overnight before centrifugation at 16.1 × 10^3^ RCF for 20 minutes to remove any unsuspended nanotube aggregates. SWCNT concentration was calculated from absorbance at 632 nm using Beer-Lambert law with extinction coefficient, ε_632_ = 0.036 L mg^-1^ cm^-1^.

### Zeta Potential Measurements of Charged SWCNTs

Zeta potential measurements were taken on a Zetasizer Nano ZS (Malvern Instruments). Prior to measurements, SWCNT samples were purified of excess salts and DNA or phospholipids by spin-filtering and diluted to a 700 μL sample at 10 mg/L concentration in MilliQ water. Three replicates of 20 measurements were obtained for each sample after 30 second equilibration.

### Streptavidin Bead Affinity Protocol on Biotin-SWCNTs

A 10 mg/L solution of SWCNT sensors was prepared in 300 μL of binding buffer (20 mM Tris-HCl, 500 mM NaCl, 1 mM EDTA, pH 7.5). 1 mg of magnetic streptavidin beads (New England Biosciences) was aliquoted into a microcentrifuge tube, a magnet was applied to the side of the tube, and the storage buffer was removed. The beads were washed 4 times by resuspension in 1000 μL of binding buffer, application of a magnet on the side of the tube, and removal of the supernatant. This wash step was repeated 3 more times. The SWCNT solution was added to the magnetic streptavidin beads and the tube was shook at a gentle speed for 2 hours. The depleted SWCNT sample was collected and the beads were washed 3 times with 1000 μL binding buffer.

### Atomic Force Microscopy of SWCNT Complexes

Monodispersed SWCNT nanosensors in the presence and absence of neutravidin were analyzed with atomic force microscopy using an Asylum Research MFP-3D AFM and TAP150AL-G-10 Silicon AFM probes (Ted Pella, tip radius < 10 nm). For the addition of neutravidin, 20 mg/L SWCNT nanosensors were incubated with 0.025 mg/mL neutravidin for 1 hour. 20 μL of SWCNT nanosensors were deposited on freshly cleaved mica and incubated for 1 hour at room temperature. Unbound nanosensors, protein, and salts were washed from the mica three times using MilliQ water. AFM was performed at a scan rate of 0.8 Hz using a sample rate of 512 lines.

## Acknowledgements

M.P.L. acknowledges support of a Burroughs Wellcome Fund Career Award at the Scientific Interface (CASI), a Stanley Fahn PDF Junior Faculty Grant with Award # PF-JFA-1760, a Beckman Foundation Young Investigator Award, a DARPA Young Faculty Award, an FFAR New Innovator Award, and a USDA award. M. P. L. is a Chan Zuckerberg Biohub Investigator and an Innovative Genomics Institute Investigator. L. C. acknowledges the support of a National Defense Science and Engineering Graduate (NDSEG) Fellowship. R. L. P. acknowledges the support of an NSF Graduate Research Fellowship. N. S. G. acknowledges the support of a FFAR Research Fellowship. Atomic force microscopy was performed at the Molecular Foundry and was supported by the Office of Science, Office of Basic Energy Sciences, of the U.S. Department of Energy under Contract No. DE-AC02-05CH11231. The authors would like to thank Darwin Yang, Tanya Chaudhury, and Jackson Travis Del Bonis-O’Donnell for insightful conversation in experimental design, as well as Eugene Kim and Jeffrey R. Long for help obtaining FTIR spectra.

## Supporting Information

Absorbance spectra and additional fluorescence spectra of SWCNTs, XPS spectra, FTIR spectra, experimental details of methanol-exchange protocol, additional zeta potential measurements

## Notes

### Competing Interest Statement

The authors have declared no competing interest.

### Summary of Updates

Materials and methods have been updated as follows (from 1 mg to 100 mg): Synthesis of SH-SWCNTs or Charged SWCNTs (NH2-SWCNTs and COOH-SWCNTs): Trz-H-SWCNTs (10 mg) were dispersed in dimethylformamide (DMF) (5 mL) and bath sonicated for 15 min at room temperature. Next, 100 mg of either cysteine, ethylenediamine, or glycine (for SH-SWCNTs, NH2-SWCNTs, and COOH-SWCNTs, respectively) and a 1.5 m excess of triethylamine to chemical were added to the mixture that was stirred at 65 C for 2 days.

## References

(1) Doane, T. L.; Burda, C. The Unique Role of Nanoparticles in Nanomedicine: Imaging, Drug Delivery and Therapy. Chem. Soc. Rev. 2012, 41 (7), 2885–2911.

(2) Scheinberg, D. A.; Grimm, J.; Heller, D. A.; Stater, E. P.; Bradbury, M.; McDevitt, M. R. Advances in the Clinical Translation of Nanotechnology. Curr. Opin. Biotechnol. 2017, 46, 66–73.

(3) Demirer, G. S.; Zhang, H.; Matos, J. L.; Goh, N.; Cunningham, F.; Sung, Y.; Chang, R.; Aditham, A. J.; Chio, L.; Cho, M.; et al. High Aspect Ratio Nanomaterials Enable Delivery of Functional Genetic Material Without DNA Integration in Mature Plants. Nat. Nanotechnol. 2019.

(4) Del Bonis-O’Donnell, J. T.; Chio, L.; Dorlhiac, G. F.; McFarlane, I. R.; Landry, M. P. Advances in Nanomaterials for Brain Microscopy. Nano Res. 2018, 11 (10), 1–29.

(5) Iverson, N. M.; Barone, P. W.; Shandell, M. A.; Trudel, L. J.; Sen, S.; Sen, F.; Ivanov, V.; Atolia, E.; Farias, E.; Mcnicholas, T. P.; et al. In Vivo Biosensing via Tissue-Localizable near-Infrared-Fluorescent Single-Walled Carbon Nanotubes. Nat. Nanotechnol. 2013, 8 (11), 873–880.

(6) Williams, R. M.; Lee, C.; Galassi, T. V.; Harvey, J. D.; Leicher, R.; Sirenko, M.; Dorso, M. A.; Shah, J.; Olvera, N.; Dao, F.; et al. Noninvasive Ovarian Cancer Biomarker Detection via an Optical Nanosensor Implant. Sci. Adv. 2018, 4 (4), eaaq1090.

(7) Kwon, O. S.; Park, S. J.; Jang, J. A High-Performance VEGF Aptamer Functionalized Polypyrrole Nanotube Biosensor. Biomaterials 2010, 31 (17), 4740–4747.

(8) Meng, L.; Zhang, X.; Lu, Q.; Fei, Z.; Dyson, P. J. Single Walled Carbon Nanotubes as Drug Delivery Vehicles: Targeting Doxorubicin to Tumors. Biomaterials 2012, 33 (6), 1689–1698.

(9) Chen, J.; Chen, S.; Zhao, X.; Kuznetsova, L. V.; Wong, S. S.; Ojima, I. Functionalized Single-Walled Carbon Nanotubes as Rationally Designed Vehicles for Tumor-Targeted Drug Delivery. J. Am. Chem. Soc. 2008, 130 (49), 16778–16785.

(10) Peng, X.; Wong, S. S. Functional Covalent Chemistry of Carbon Nanotube Surfaces. Adv. Mater. 2009, 21 (6), 625–642.

(11) Hilderbrand, S. A.; Weissleder, R. Near-Infrared Fluorescence: Application to in Vivo Molecular Imaging. Curr. Opin. Chem. Biol. 2010, 14 (1), 71–79.

(12) Bonis-O’Donnell, J. T. D.; Page, R. H.; Beyene, A. G.; Tindall, E. G.; McFarlane, I. R.; Landry, M. P. Dual Near-Infrared Two-Photon Microscopy for Deep-Tissue Dopamine Nanosensor Imaging. Adv. Funct. Mater. 2017, 27 (39), 1–10.

(13) Bachilo, S. M.; Strano, M. S.; Kittrell, C.; Hauge, R. H.; Smalley, R. E.; Weisman, R. B. Structure-Assigned Optical Spectra of Single-Walled Carbon Nanotubes. Science (80-.). 2002, 298 (5602), 2361–2366.

(14) Kruss, S.; Landry, M. P.; Vander Ende, E.; Lima, B.; Reuel, N. F.; Zhang, J.; Nelson, J. T.; Mu, B.; Hilmer, A. J.; Strano, M. S. Neurotransmitter Detection Using Corona Phase Molecular Recognition on Fluorescent Single-Walled Carbon Nanotube Sensors. J. Am. Chem. Soc. 2014, 136 (2), 713–724.

(15) Zhang, J.; Landry, M. P.; Barone, P. W.; Kim, J.-H.; Lin, S.; Ulissi, Z. W.; Lin, D.; Mu, B.; Boghossian, A. A.; Hilmer, A. J.; et al. Molecular Recognition Using Corona Phase Complexes Made of Synthetic Polymers Adsorbed on Carbon Nanotubes. Nat. Nanotechnol. 2013, 8 (12), 959–968.

(16) Landry, M. P.; Ando, H.; Chen, A. Y.; Cao, J.; Kottadiel, V. I.; Chio, L.; Yang, D.; Dong, J.; Lu, T. K.; Strano, M. S. Single-Molecule Detection of Protein Efflux from Microorganisms Using Fluorescent Single-Walled Carbon Nanotube Sensor Arrays. Nat. Nanotechnol. 2017, 12, 368–377.

(17) Bisker, G.; Dong, J.; Park, H. D.; Iverson, N. M.; Ahn, J.; Nelson, J. T.; Landry, M. P.; Kruss, S.; Strano, M. S. Protein-Targeted Corona Phase Molecular Recognition. Nat. Commun. 2016, 7, 10241.

(18) Bisker, G.; Bakh, N. A.; Lee, M. A.; Ahn, J.; Park, M.; O’Connell, E. B.; Iverson, N. M.; Strano, M. S. Insulin Detection Using a Corona Phase Molecular Recognition Site on Single-Walled Carbon Nanotubes. ACS Sensors 2018, 3 (2), 367–377.

(19) Chio, L.; Del Bonis-O’Donnell, J. T.; Kline, M. A.; Kim, J. H.; McFarlane, I. R.; Zuckermann, R. N.; Landry, M. P. Electrostatic Assemblies of Single-Walled Carbon Nanotubes and Sequence-Tunable Peptoid Polymers Detect a Lectin Protein and Its Target Sugars. Nano Lett. 2019, 19 (11), 7563–7572.

(20) Hong, G.; Diao, S.; Chang, J.; Antaris, A. L.; Chen, C.; Zhang, B.; Zhao, S.; Atochin, D. N.; Huang, P. L.; Andreasson, K. I.; et al. Through-Skull Fluorescence Imaging of the Brain in a New near-Infrared Window. Nat. Photonics 2014, 8 (9), 723–730.

(21) Harvey, J. D.; Jena, P. V.; Baker, H. A.; Zerze, G. H.; Williams, R. M.; Galassi, T. V.; Roxbury, D.; Mittal, J.; Heller, D. A. A Carbon Nanotube Reporter of MicroRNA Hybridization Events in Vivo. Nat. Biomed. Eng. 2017, 1(March), 0041.

(22) Beyene, A. G.; Delevich, K.; Del Bonis-O’Donnell, J. T.; Piekarski, D. J.; Lin, W. C.; Wren Thomas, A.; Yang, S. J.; Kosillo, P.; Yang, D.; Prounis, G. S.; et al. Imaging Striatal Dopamine Release Using a Nongenetically Encoded near Infrared Fluorescent Catecholamine Nanosensor. Sci. Adv. 2019, 5 (7), 1–12.

(23) Freeley, M.; Worthy, H. L.; Ahmed, R.; Bowen, B.; Watkins, D.; Macdonald, J. E.; Zheng, M.; Jones, D. D.; Palma, M. Site-Specific One-to-One Click Coupling of Single Proteins to Individual Carbon Nanotubes: A Single-Molecule Approach. J. Am. Chem. Soc. 2017, 139 (49), 17834–17840.

(24) Banerjee, S.; Hemraj-Benny, T.; Wong, S. S. Covalent Surface Chemistry of Single-Walled Carbon Nanotubes. Adv. Mater. 2005, 17 (1), 17–29.

(25) Onitsuka, H.; Fujigaya, T.; Nakashima, N.; Shiraki, T. Control of the Near Infrared Photoluminescence of Locally Functionalized Single-Walled Carbon Nanotubes via Doping by Azacrown-Ether Modification. Chem. - A Eur. J. 2018, 24 (37), 9393–9398.

(26) Kwon, H.; Furmanchuk, A.; Kim, M.; Meany, B.; Guo, Y.; Schatz, G. C.; Wang, Y. Molecularly Tunable Fluorescent Quantum Defects. J. Am. Chem. Soc. 2016, 138 (21), 6878–6885.

(27) Palma, M.; Wang, W.; Penzo, E.; Brathwaite, J.; Zheng, M.; Hone, J.; Nuckolls, C.; Wind, S. J. Controlled Formation of Carbon Nanotube Junctions via Linker-Induced Assembly in Aqueous Solution. J. Am. Chem. Soc. 2013, 135 (23), 8440–8443.

(28) Luo, H. Bin; Wang, P.; Wu, X.; Qu, H.; Ren, X.; Wang, Y.One-Pot, Large-Scale Synthesis of Organic Color Center-Tailored Semiconducting Carbon Nanotubes. ACS Nano 2019, 13 (7), 8417–8424.

(29) Setaro, A.; Adeli, M.; Glaeske, M.; Przyrembel, D.; Bisswanger, T.; Gordeev, G.; Maschietto, F.; Faghani, A.; Paulus, B.; Weinelt, M.; et al. Preserving π-Conjugation in Covalently Functionalized Carbon Nanotubes for Optoelectronic Applications. Nat. Commun. 2017, 8, 1–7.

(30) Godin, A. G.; Setaro, A.; Gandil, M.; Haag, R.; Adeli, M.; Reich, S.; Cognet, L. Photoswitchable Single-Walled Carbon Nanotubes for Super-Resolution Microscopy in the near-Infrared. Sci. Adv. 2019, 5 (9), eaax1166.

(31) Beyene, A. G.; Alizadehmojarad, A. A.; Dorlhiac, G.; Goh, N.; Streets, A. M.; Král, P.; Vuković, L.; Landry, M. P. Ultralarge Modulation of Fluorescence by Neuromodulators in Carbon Nanotubes Functionalized with Self-Assembled Oligonucleotide Rings. Nano Lett. 2018, 18 (11), 6995–7003.

(32) Carr, J. A.; Franke, D.; Caram, J. R.; Perkinson, C. F.; Saif, M.; Askoxylakis, V.; Datta, M.; Fukumura, D.; Jain, R. K.; Bawendi, M. G.; et al. Shortwave Infrared Fluorescence Imaging with the Clinically Approved Near-Infrared Dye Indocyanine Green. Proc. Natl. Acad. Sci. 2018, 201718917.

(33) Wang, H.; Michielssens, S.; Moors, S. L. C.; Ceulemans, A. Molecular Dynamics Study of Dipalmitoylphosphatidylcholine Lipid Layer Self-Assembly onto a Single-Walled Carbon Nanotube. Nano Res. 2009, 2 (12), 945–954.

(34) Lee, H.; Kim, H. Self-Assembly of Lipids and Single-Walled Carbon Nanotubes: Effects of Lipid Structure and PEGylation. J. Phys. Chem. C 2012, 116 (16), 9327–9333.

(35) Streit, J. K.; Fagan, J. A.; Zheng, M. A Low Energy Route to DNA-Wrapped Carbon Nanotubes via Replacement of Bile Salt Surfactants. Anal. Chem. 2017, 89 (19), 10496–10503.

(36) Giraldo, J. P.; Landry, M. P.; Kwak, S.-Y.; Jain, R. M.; Wong, M. H.; Iverson, N. M.; Ben-Naim, M.; Strano, M. S. A Ratiometric Sensor Using Single Chirality Near-Infrared Fluorescent Carbon Nanotubes: Application to in Vivo Monitoring. Small 2015, 11 (32), 3973–3984.

(37) Fröhlich, E. The Role of Surface Charge in Cellular Uptake and Cytotoxicity of Medical Nanoparticles. Int. J. Nanomedicine 2012, 7, 5577–5591.

(38) Mann, F. A.; Lv, Z.; Grosshans, J.; Opazo, F.; Kruss, S. Nanobody Conjugated Nanotubes for Targeted Near-Infrared in Vivo Imaging and Sensing. Angew. Chemie Int. Ed. 2019.

(39) Marttila, A. T.; Laitinen, O. H.; Airenne, K. J.; Kulik, T.; Bayer, E. A.; Wilchek, M.; Kulomaa, M. S. Recombinant NeutraLite Avidin: A Non-Glycosylated, Acidic Mutant of Chicken Avidin That Exhibits High Affinity for Biotin and Low Non-Specific Binding Properties. FEBS Lett. 2000, 467 (1), 31–36.

(40) Neish, C. S.; Martin, I. L.; Henderson, R. M.; Edwardson, J. M. Direct Visualization of Ligand-Protein Interactions Using Atomic Force Microscopy. Br. J. Pharmacol. 2002, 135 (8), 1943–1950.

(41) Kundu, S.; Wang, Y.; Xia, W.; Muhler, M.Thermal Stability and Reducibility of Oxygen-Containing Functional Groups on Multiwalled Carbon Nanotube Surfaces : A Quantitative High-Resolution XPS and TPD / TPR Study. 2008, 16869–16878.

